# Genotyping by sequencing of 393 *Sorghum bicolor* BTx623 × IS3620C recombinant inbred lines improves sensitivity and resolution of QTL detection

**DOI:** 10.1101/308478

**Authors:** WenQian Kong, Changsoo Kim, Dong Zhang, Hui Guo, Xu Tan, Huizhe Jin, Chengbo Zhou, Lan-shuan Shuang, Valorie Goff, Uzay Sezen, Gary Pierce, Rosana Compton, Cornelia Lemke, Jon Robertson, Lisa Rainville, Susan Auckland, Andrew H. Paterson

## Abstract

We describe a genetic map with a total of 381 bins of 616 genotyping by sequencing (GBS)-based SNP markers in a F_6_-F_8_ recombinant inbred line (RIL) population of 393 individuals derived from crossing *S. bicolor* BTx623 to *S. bicolor* IS3620C, a guinea line substantially diverged from BTx623. Five segregation distorted regions were found with four showing enrichment for *S. bicolor* alleles, suggesting possible selection during formation of this RIL population. A quantitative trait locus (QTL) study with this number of individuals, tripled relative to prior studies of this cross, provided resources, validated previous findings, and demonstrated improved power to detect plant height and flowering time related QTLs relative to other published studies. An unexpected low correlation between flowering time and plant height permitted us to separate QTLs for each trait and provide evidence against pleiotropy. Ten non-random syntenic regions conferring QTLs for the same trait suggest that those QTLs may represent alleles at genes functioning in the same manner since the 96 million year ago genome duplication that created these syntenic relationships, while syntenic regions conferring QTLs for different trait may suggest sub-functionalization after duplication. Collectively, this study provides resources for marker-assisted breeding, as well as a framework for fine mapping and subsequent cloning of major genes for important traits such as plant height and flowering time in sorghum.

## Introduction

The most drought resistant of the world’s top five cereal crops, sorghum, contributes 26-29 percent of calories in the human diet in semi-arid areas of Africa (FAO), on some of the world’s most degraded soils and often with limited water inputs. A multi-purpose crop, sorghum has been traditionally used as grain and straw, and is also a promising crop for bioenergy production from starch, sugar, or cellulose on marginal lands with limited water and other resources (Rooney et al. 2007). Botanically, *Sorghum bicolor* is a model for plants that use C4 photosynthesis, improving carbon assimilation especially at high temperature, complementary to C3 model plants such as *Oryza sativa*. The sequenced ~730 megabase sorghum genome (Paterson et al. 2009) has not experienced genome duplication in an estimated ~96 million years (Wang et al. 2015), making it particularly useful to study other C4 plants with large polyploid genomes such as maize and many other grasses in the *Saccharinae* clade including Miscanthus and Saccharum (sugarcane).

Evolution, natural selection and human improvement of sorghum have contributed to great morphological diversity of *Sorghum bicolor* spp. Cultivated forms of this species can be classified into five botanical races, bicolor, guinea, caudatum, kafir and durra; with ten intermediate races recognized based on inflorescence architecture and seed morphology (Dewet and Huckabay 1967; Harlan and Dewet 1972). *Sorghum bicolor* also has many wild relatives such as *S. propinquum* (2n=2x=20), diverged from *S. bicolor* 1-2 million years ago (Feltus et al. 2004); *Sorghum halepense* (2n=4x=40), an invasive and weedy species formed by unintentional crossing of *S. bicolor* and *S. propinquum* (Paterson et al. 1995a); and many hybrids between these species. These many possible intra- and interspecific crosses have made sorghum a particular interesting model for dissecting the genetic control of complex traits such as plant height and maturity (Lin et al. 1995; Hart et al. 2001; Feltus et al. 2006; Takai et al. 2012), tillering and vegetative branching (Alam et al. 2014; Kong et al. 2014; Kong et al. 2015), perenniality related traits (Paterson et al. 1995b; Washburn et al. 2013), sugar composition (Murray et al. 2008a; Murray et al. 2008b; Ritter et al. 2008; Shiringani et al. 2010; Vandenbrink et al. 2013), stay-green leaves and plants (Haussmann et al. 2002; Kassahun et al. 2010), drought resistance (Tuinstra et al. 1998; Kebede et al. 2001; Sanchez et al. 2002), disease and insect resistance (Katsar et al. 2002; Totad et al. 2005).

*Sorghum bicolor* IS3620C is a representative of botanical race ‘guinea’, and is substantially diverged from *S. bicolor* BTx623 among the sorghum races. Prior workers described a genetic map with 323 RFLP and 147 SSR markers (Bhattramakki et al. 2000; Kong et al. 2000), and a quantitative trait locus (QTL) study with 137 F6-8 RILs demonstrated its usefulness for QTL detection by discovering as many as 27 QTLs traits of agronomical importance, such as plant height, maturity, number of basal tillers and panicle length (Hart et al. 2001; Feltus et al. 2006).

Next generation sequencing (NGS) has brought new power to revealing allelic differences among individuals by detecting large numbers of single nucleotide polymorphisms (SNP). While it is neither necessary nor practical (yet) to generate whole genome sequences for every individual in a designed mapping population, reduced representation library (RRL) sequencing has been widely utilized and proved to be both a cost and labor efficient tool for genotyping (Andolfatto et al. 2011a; Kim et al.). Although current genotyping by sequencing (GBS) platforms and pipelines still face various issues regarding accuracy (especially in polyploids with high levels of heterozygosity), GBS is still a very powerful tool to generate genetic maps with high quality and resolution in nearly homozygous populations with reference genome sequences available (Kim et al. 2015).

In this paper, we describe a genetic map with a total of 381 bins of 616 GBS-based SNP markers in a recombinant inbred line (RIL) population of 393 individuals derived from two divergent *S. bicolor* genotypes, BTx623 and IS3620C. Recent work that utilized digital genotyping to saturate the genetic map of 137 individuals (Morishige et al. 2013) has validated QTLs for height more precisely but showed little increase in power to detect QTLs relative to prior analysis using a lower density of markers (Hart et al. 2001). Our study has tripled the number of individuals described in previous studies (Bhattramakki et al. 2000; Kong et al. 2000; Hart et al. 2001; Morishige et al. 2013), increasing both the number of discernable recombination events and the number of individuals carrying each parental allele, increasing power to detect QTLs. A quantitative trait analysis of days to flowering and components of plant height demonstrate the improved power to detect quantitative trait loci (QTL) in this expanded population. Syntenic relationships among sorghum QTLs may reflect homeologous regions retaining the same or similar function after a genome duplication 96 million years ago (Wang et al. 2015). Results of this new QTL mapping build on those of many other studies (Lin et al. 1995; Hart et al. 2001; Kebede et al. 2001; Brown et al. 2006; Srinivas et al. 2009; Shiringani et al. 2010; Morris et al. 2013; Zhang et al. 2015), enriching current resources and providing a better understanding of the genetic control of plant height and days to flowering of *S. bicolor*.

## Materials and Methods

### Genetic stocks

The mapping population is comprised of 399 F7-8 RILs derived by selfing a single F_1_ plant from *S. bicolor* BTx623 and IS3620C as described (Kong et al. 2000; Hart et al. 2001; Burow et al. 2011). This RIL population was planted at the University of Georgia Plant Science Farm, Watkinsville, GA, USA on 10 May 2011 and 18 May 2012. Single 3-m plots of each RIL were machine planted in a completely randomized design. Two representative samples were phenotyped for each genotype.

### Genotyping

Leaf samples of the RIL population was frozen at −80 C and lyophilized for 48 hours. Genomic DNA was extracted from the lyophilized leaf sample based on Aljanabi et al. (1999).

Our GBS platform is a slightly modified version of Multiplex Shotgun Genotyping (MSG) (Andolfatto et al. 2011b) combined with the Tassel GBS analysis pipeline. The basic sequencing platform is an in-house Illumina MiSeq that generates up to 25 million reads of 150 base pairs (bps) fragments per run with single-end sequencing. With one sequencing run of this platform, we obtained 7103 raw SNPs and 691 polymorphic SNPs that were sufficient for genetic mapping for this RIL population (Kim et al. 2015).

SNP ‘calling’ (inference) was based on the reference genome sequence of *Sorghum bicolor* (Paterson et al. 2009). Alignment used the Burrows-Wheeler Aligner (BWA) for single end read (samse). In TASSEL-GBS, the first 64 bps of each reads were mapped onto a reference genome to decide the position of the reads. SNPs were called based on the alignment of reads to the reference genome. Heterozygosity at a locus is called if two alleles are each inferred to be present at a probability greater than that of sequencing error. Raw SNP data from the Tassel GBS pipeline were further filtered based on several criteria: (a) SNPs were removed if the minor allele frequency is less than 5% or the proportion of missing genotypes greater than 40%; (b) In order to reduce the number of redundant SNPs in studies where strong linkage disequilibrium necessitates only 5–10 cM resolution, we merged SNPs for which pairwise linkage disequilibrium (r^2^) is greater than 0.9, deriving consensus genotypes in a manner minimizing missing genotypes. SNPs are further merged if the Pearson’s correlation between them is larger than 0.95; (c) For bi-parental populations, the missing genotype of one parental line can be imputed by offspring genotypes if the genotype of the other parent at the locus is known. After these filtering steps, SNP data are used for genetic mapping.

### Map construction

A genetic map using 616 SNP markers was created using R/qtl (Broman et al. 2003). We further assigned bins for each chromosome to merge markers within 1 cM in genetic distance. Bin genotypes were defined as follows: If there was only one marker in the bin, the bin genotype would be the same as the marker genotype; if there were more than one marker in the bin, bin genotypes would be determined by merging marker genotypes to minimize missing data points. For example, for a particular individual if there were three SNP markers in a bin, and if the marker genotypes for all three SNPs agree, the bin genotype will be the same as the marker genotypes; if the marker genotypes showed discrepancy but not due to missing data, the bin genotype would be missing data, if more than one genotype were missing, the bin genotype would be the same as the non-missing genotype. Following this method, we obtained a total of 381 bins for map construction. Marker ordering used both *de novo* and reference based methods, i.e., the physical positions of SNPs. The ‘ripple’ function was used to assist and validate ordering of the genetic map.

We used a chi-squared test to calculate the deviation from expected ratio (1:1) for each marker with both raw and imputed data as an indicator for segregation distortion. To account for multiple comparisons across the genome, the significance level was adjusted using Bonferroni correction. The imputed data was generated using R/qtl (Broman et al. 2003).

### QTL mapping

QTLs were detected for five traits of interest: plant height (**PH**), the overall length of a plant; base to flag length (**BTF**), the length from the base of the plant to the flag leaf; flag to rachis length (**FTR**), the length from the flag leaf to rachis (a positive sign was assigned if the position of the rachis is taller than the flag leaf; a negative value was assigned if the rachis was ‘buried’ in the flag leaf); number of nodes (**ND**); and days to flowering (**FL**), the average days to flowering for the first five plants for each genotype.

We combined the phenotypic data using Best Linear Unbiased Prediction (BLUP) by treating individuals, years, replications nested within years and the interactions between individuals and years as random, since heritability for the traits of interest were relatively high. In 2011, we observed and recorded a soil type change within the experimental fields, which was treated as a covariate to calculate BLUP values for each genotype. A genome scan with the interval mapping method was first conducted with 1000 permutation tests; the putative QTLs were then selected and fit into a multiple QTL model. We added additional QTLs to the model if they exceeded the threshold of 3.0 after fixing the effect of QTLs included in the first genome scan. A multiple QTL model (MQM) was used to determine the final model for each trait. All statistical analyses and QTL mapping used R (R Core Team 2016) and the R/QTL package (Broman et al. 2003).

QTL nomenclature used a system that was previously described in rice (McCouch et al. 1997), starting with a ‘q’, followed by an abbreviation for each trait (**PH, BTF, FTR, ND** and **FL**), then the chromosome number and a decimal number to differentiate multiple QTLs on the same chromosome.

### Data availability

File S1 contains genotypes for the bin map. File S2 contains genotypes for the original map. File S3 contains the genomic positional information for bin markers. File S4 and S5 contain phenotypes from 2011 and 2012 respectively.

## Results

### Genetic map

A total of 399 RILs were genotyped with 690 SNP markers. Six individuals for which genotyping data suggested three times more than the average number of recombination events were deemed erroneous and removed from the analysis. Marker ordering first follows the published sorghum genome sequence (Paterson et al. 2009). A *de novo* marker ordering method is also used to compare the order of the genetic map with the reference-based method, but no obvious differences were observed for these two methods in terms of the LOD scores. We excluded 74 unlinked SNPs and obtained an initial genetic map with a total of 616 markers on the ten sorghum chromosomes. As detailed in the methods, we combined SNPs that are within 1cM in genetic distance to construct a genetic map with 381 bins (bin map) with varying SNP numbers in each bin (File S3). The bin map collectively spans a genetic distance of 1404.8 cM, with average spacing of 3.8 cM between loci and the largest gap being 27.6 cM on chromosome 5 (Table 1). The percentages of missing genotypes are 24% and 18.4% for the initial and bin maps, respectively. About 54.2% of the alleles of the RIL population come from *S. bicolor* BTx623, and 45.8% from *S. bicolor* IS3620C.

The order of the genetic map agrees closely with the physical positions of loci on the genome sequence (Figure 1), suggesting that the GBS method used yields a high quality genetic map in this nearly-homozygous diploid population. Markers are generally more concentrated in the distal regions of each chromosome than central regions (consistent with the general distribution of low-copy DNA sequences in sorghum: Paterson et al 2008), although distributions of markers on each chromosome vary. For example, we have observed a much larger pericentrimeric region on chromosome 7 than on chromosome 1.

### Segregation distortion

Segregation distortion occurs when the segregation ratio of offspring at a locus deviates from the Mendelian expectation. In a RIL population, we expect to see half of the alleles come from each parent, i.e., the expected segregation ratio is 1:1 for each marker locus. A deviation from this ratio may be a result of gametic or zygotic selection. The BTx623 × IS3620C genetic map reveals several clusters of markers experiencing segregation distortion on chromosomes 1, 4, 5, 8, and 9 (Figure 2), peaking at 48.0, 153.6, 64.4, 47.5, and 122.4 cM, corresponding to 14.8, 65.1, 49.9, 27.8-38.5 and 59.5 in physical distance, respectively, with the imputed data. All regions but the one on chromosome 8 show enrichment of *S. bicolor* alleles. The most extreme segregation distorted region is on chromosome 1, spanning 0-105 cM (Figure 2) with a ratio of 358:35 (p = 1.10E-59) at the peak marker, S1_14765342. Peng et al. (1999) also observed this long-spanning segregation distorted region on chromosome 1, despite using a population with a much smaller sample size and marker numbers.

Interestingly, the same general region of extreme segregation distortion on chromosome 1 is also found in two *S. bicolor* × *S. propinquum* derived populations (Bowers et al. 2003; Kong 2013), and remarkably, all three populations mapped this distortion peak at ~14 Mb in physical distance. This correspondence between populations suggests that alleles from *S. bicolor* might be selected for at this genomic position, however, the exact mechanism and the genes involved are unclear.

### QTL Mapping

The present BTx623 × IS3620C RIL map provides higher power for detecting QTLs than a prior map of a subset of these progenies. As examples, we have investigated QTLs for five phenotypic traits, plant height (**PH**), base to flag length (**BTF**), flag to rachis length (**FTR**), number of nodes (**ND**) and days to flowering (**FL**). Means, standard deviation and other summary statistics are shown in Table S1. Broad-sense heritability estimates for all five traits are relatively high (Table S1). It is interesting that the average **PH** of the progenies is 98.62 cm, greater than the average of either parents, 94.40 for BTx623 and 84.49 for IS3620C. Likewise, **BTF is** greater than the average of either parent. Both height components show substantial genetic variation, indicating that each parent contributes different alleles for **PH** to their progenies (Table S1). Moreover, the relatively high genetic variation in **PH** in this population fosters discovery of QTLs, despite that the difference between the two parents is relatively small.

**PH** and **BTF** are highly correlated in this population (Table S2), with a correlation coefficient of 0.8983 (p<0.001). We detect a total of seven and five QTLs for **PH** and **BTF**, accounting for 40.13% and 41.58% of the total phenotypic variances, respectively (Table S3). Three QTLs on chromosomes 3, 6 and 7 overlap for these two traits. The QTLs on chromosomes 6 and 7 account for the majority of the phenotypic variance explained for these traits, ~28% and ~32% for **PH** and **BTF**, respectively. These two large effect QTLs might be related to previously defined **PH** genes, presumably *dw2* on chromosome 6 and *dw3* (Sb07g023730) on chromosome 7 (Quinby and Karper 1945; Multani et al. 2003).

Not only do we detect more QTLs than were found in the previous study of a subset of this population (Hart et al. (2001)), seven QTLs for **PH** and nine QTLs for **FL** in our study versus five and three from the previous study, but some of the QTL intervals also significantly reduced (Table S3). For example, the 1-lod interval of QTL on chromosome 7 is narrowed from ~26 cM previously to only ~3 cM in this study (from 57.7 Mb to 59.5 Mb in physical distance), and harbors the gene Sb07g023730 (*DW3*) at ~58.6 Mb. This example indicates that nearly tripling the numbers of individuals and increasing marker density greatly increased the power of QTL detection.

An interesting phenotype that we observed is distance from the flag leaf to the rachis. We distinguished whether or not the rachis is immersed in the flag leaf (see Materials and Methods). The correlation coefficient (Table S2) between **FTR** and **PH** (r=0.0778), though significant at p<0.01, is not nearly high as the correlation between the **BTF** and **PH** (r=0.8983), suggesting that the genetic control of these two traits might be different. Indeed, QTL mapping suggests that the genetic control of **FTR** is quite different from that of **PH** and **BTF**, with only one QTL (qFTR7.1) on chromosome 7 overlapping with QTL for **PH** and **BTF** (qPH7.1 and qBTF7.1). While the one-LOD QTL interval for qPH10.1 overlaps with qFTR10.1 to some extent, there is no solid evidence to conclude that they are controlled by the same genetic factors, given that the likelihood peaks of the QTLs for these two traits are ~10 cM apart. We detected a total of five QTLs for **FTR**, explaining 28.21 % of the total phenotypic variance (Table S3). The additive effect of **FTR** needs to be carefully interpreted, especially when compared to the additive effect of **PH**. Since the average value of **FTR** is negative, a negative number for the additive effect for this trait indicates increased **FTR** for a particular allele. For example, the additive effect for qFTR7.1 is −1.19, which indicates that FTR of plants carrying the IS3620 alleles are actually longer than those of plants carrying the BTx623 alleles. Both alleles from IS3620C for qPH7.1 and qFTR7.1 have the same effect of increasing length, although the sign of their additive effects is different.

We have detected a total of six QTLs for number of nodes (**ND**) in this population (Figure 3), collectively explaining 32.07% of the phenotypic variance. The largest effect QTL is qND8.1, with a LOD score of 12.96 and explaining 11.15% of the variance. QTLs for **ND** rarely overlap with other **PH** related traits—only qND10.1 shows some correspondence with other height related traits, marginally overlapping with qFTR10.1.

In this study, **PH** and **FL** were not significantly correlated (Table S2), an unexpected finding compared to many other sorghum studies (Lin et al. 1995; Murray et al. 2008b; Ritter et al. 2008). A total of nine QTLs are detected for **FL**, substantially more than the 4-6 conventionally thought to influence this trait in a wide range of sorghum genotypes (Quinby and Karper 1945), although only collectively explaining 46.33% of the total phenotypic variance. QTL intervals for **FL** rarely overlap with those for **PH**, **BTF** and **FTR**. However, five out of six QTLs for **ND** overlap with QTLs for **FL**, and the sign of the allelic effect suggests that plants with more nodes usually flower late. This result indicates that some genes might have pleiotropic effects on these traits or genes for that these two traits are linked and have been selected simultaneously.

### QTL correspondence with other studies

Traits related to sorghum **PH** and **FL** have been extensively studied in many QTL experiments (Lin et al. 1995; Hart et al. 2001; Kebede et al. 2001; Brown et al. 2006; Ritter et al. 2008; Srinivas et al. 2009; Shiringani et al. 2010) and two genome wide association studies (Morris et al. 2013; Kong et al. 2015). The Comparative Saccharinae Genome Resource QTL database (Zhang et al. 2013) aids comparisons of QTL intervals across different studies in sorghum and facilitates validation of QTLs for traits of interest. All our QTLs detected for **PH** have been found in other studies. Three **PH** QTLs, qPH6.1, qPH7.1 and qPH9.1, also found in two GWAS studies (Morris et al. 2013; Zhang et al. 2015), are likely to correspond to three dwarfing genes in sorghum, *dw2, dw3* and *dw1* (Quinby and Karper 1945). The accuracy of our QTL study can be demonstrated by the gene known to cause the *dw3* phenotype, Sb07g023730, on chromosome 7. Our QTL study narrowed the 1-lod interval for this QTL to 2Mb (57.7-59.5Mb), with a peak at 58.4Mb, close to the 58.6 Mb location of the causal gene (Multani et al. 2003).

A total of 6 QTLs controlling **FL** in the BTx623 × IS3620C RILs, qFL1.2, qFL3.1, qFL6.1, qFL8.2, qFL9.1 and qFL10.1, showed correspondence with QTLs found in other studies, (Lin et al. 1995; Hart et al. 2001; Brown et al. 2006; Shiringani et al. 2010; Yang et al. 2014) and two QTLs, qFL1.1 and qFL6.1, are novel. The QTLs with the largest effects on **FL**, qFL8.2 with a LOD score of 16.1 and explaining 11.16% of the phenotypic variance; and qFL9.1 with a LOD score of 11.4 and explaining 7.69% of phenotypic variance, have been consistently found in many independent QTL and GWAS studies (Lin et al. 1995; Brown et al. 2008; Morris et al. 2013; Zhang et al. 2015). Identification of the genes underlying these QTL regions might be especially important. The peak of qFL9.1 is located at ~59.3Mb in our study, close to significant peaks at ~58.7Mb and ~59.0Mb for **FL** found in a GWAS study (Zhang et al. 2015).

### Syntenic study

A total of 30 out of 202 genomic regions contain QTLs found in this study were located in colinear locations within sorghum (Paterson et al. 2009) resulting from genome duplication events (Table 2 and Figure 4), as confirmed using the Plant Genome Duplication Database (Lee et al. 2013). Among these, a total of five regions on chromosomes 1 (1), 3 (3), 9 (1) are duplicated within the same chromosome. Among the 25 duplicated genomic regions located on different chromosomes, ten syntenic regions contain the same trait (2 for ND, 3 for PH, 4 for FL and 1 for BTF), which is significantly more than expected to occur by chance (p=0.0002), and 15 regions contain different traits (Figure 4).

## Discussion

The present study providing a relatively dense GBS-based map for 393 individuals of the singularly-important *S. bicolor* BTx623×IS3620C RIL population identifies new QTLs and increases precision of mapping previously-known QTLs, providing both an important resource and new information about the genetic control of important sorghum traits. Benefiting both from an increased sample size and GBS, our study has demonstrated increased power and accuracy of detecting QTLs, relative to previous studies of a total of 137 individuals (Hart et al. 2001; Feltus et al. 2006). We also discovered a total of five regions with segregation distortion, possibly due to gametic or zygotic selection during the formation of this RIL population.

There are several advantages of the bin mapping strategy used in this paper. First, it reduces the percentage of missing genotypes by combining the genotypes of adjacent markers. For this experiment, the percentage of missing genotypes is reduced by 5.6% with the bin map. Moreover, QTL intervals are usually 5-10 cM for a typical bi-parental QTL experiment; high marker density does not significantly increase the power of detecting QTLs (Lander and Botstein 1989) while combining markers may increase computational processing speed.

Within this single population, we now find more QTLs for **PH** and **FL** than have been classically thought to segregate in all forms of *Sorghum bicolor* (Quinby and Karper 1945), reiterating a conclusion from meta-analysis of multiple populations (Zhang et al. 2015) that these traits could not be accounted for by the 4-6 loci suggested by classical studies. Most of the QTLs we mapped here correspond to QTLs found in other studies, improving confidence in our result. An example is re-identification of the *dw3* locus on chromosome 7, a P-glycoprotein auxin transporter (Multani et al. 2003), which proved the power and accuracy of our QTL study by narrowing the QTL interval from ~26 cM to 3 cM known to harbor the causal gene. More generally, QTL mapping complements other data types toward identification of causal genes. For example, a QTL on chromosome 6, possibly the classical *dw2* locus, has been refined to a 5 cM interval in this study, with a likelihood peak at ~42.4 Mb. A GWAS study (Morris et al. 2013) proposed that *dw2* lies between 39.7 Mb-42.6 Mb. Other recent studies have proposed nearby candidate genes for *dw2* including Sobic.006G067700 (Sb06g15430, (Hilley et al. 2017)) and Sb06g007330 (Cuevas et al. 2016).

In addition to **PH** and **FL**, we have also identified QTLs for three other traits that are not extensively studied, **BTF**, **FTR** and **ND**. QTL intervals for **FTR** mostly differ from those associated with **PH**, suggesting that the genetic control of these traits might be different. We also found that the genetic control of the **ND** is correlated with **FL**, demonstrated by the fact that five of six QTLs for **ND** correspond to QTLs for **FL**.

An unexpected low correlation between **FL** and **PH** in this population and the high power of this genetic map, together permit us to differentiate **FL** and **PH** QTLs. In fact, no corresponding QTLs for these two traits were identified in the present study, an extremely unusual finding. This lack of overlap strongly supports a hypothesis (Cuevas et al. 2016(Lin et al 1995) that the *dw2* trait affecting **PH** and the *ma1* trait affecting flowering, each mapping very close together on chromosome 6, are determined by different genes. This is proven by the fact that alternative alleles at the *ma1* locus do not segregate in this study (IS3620C was ‘converted’ to day-neutral flowering, thus has *ma1* as does BTx623) (Stephens et al. 1967) while a strong signal has been detected for **PH** on chromosome 6 at ~42.4cM, in the vicinity of a recently published *dw2* candidate gene (Hilley et al. 2017), albeit further functional analysis is needed. A similar example is within the general area of *dw1* (Yamaguchi et al. 2016) on chromosome 9 (qPH9.1) where we also find a QTL controlling **FL**, qFL9.1, that is ~30cM from qPH9.1. This result again suggests that two separate QTLs control these **PH** and **FL** effects, a conclusion that is also supported by another study (Thurber et al. 2013).

We discovered a total of 30 syntenic regions within the sorghum genome sequence (Paterson et al. 2009; Lee et al. 2013) containing QTLs, with 10 regions containing QTLs responsible for the same trait (Table 2 and Figure 4). This non-random correspondence between regions of the genome conferring the same traits indicates that the ten syntenic regions contain corresponding (homoeologous) genes that may still function in the same ways despite being duplicated 96 million years ago (Wang et al. 2015), while the syntenic regions with different traits may suggest potential sub-functionalization of genes after duplication.

Components of **PH** and **FL** have been and will continue to be important for sorghum breeding programs. The past century has witnessed breeding for modern varieties with a particular plant type, for example a semi-dwarf type, to realize striking increases in production such as those which led to the “Green Revolution” (Evenson and Gollin 2003). The concept of ideotype breeding (Donald (1968), is still an ongoing priority for many breeding programs to increase food and feed production, adapt to climate change and minimize inputs. Genetic components discovered for plant height related traits and flowering time in this study, together with closely-linked diagnostic DNA markers that permit their selection at seedling stages or in non-target environments, may benefit breeding for plant types idealized for the different purposes that sorghum is used. Specifically, little correspondence between **PH** and **FL**, together with narrowed QTL intervals, facilitates accurate selection for each trait. The QTLs found in this study and their correspondence with those from many other studies also provides a framework for fine mapping or subsequent cloning of major genes for **PH** and **FL** in sorghum.

## Acknowledgments

The late K. Schertz, USDA-ARS and Texas A&M University, was instrumental in producing the BTx623×IS3620C RILs. We thank the members of the Paterson lab for valuable help and suggestions. We appreciate financial support from the AFRI Plant Growth and Development Program (2009-03477), DOE-USDA Plant Feedstock Genomics Program (2012-03304), and USDA Biotechnology Risk Assessment Program (2012-01658).

